# Effects of the Ordering of Natural Selection and Population Regulation Mechanisms on Wright-Fisher Models

**DOI:** 10.1101/061911

**Authors:** Zhangyi He, Mark Beaumont, Feng Fu

## Abstract

The Wright-Fisher model and its extensions are of central importance in population genetics, and so far, they have formed the basis of most theoretical and applied population genetic research. In the present work, we explore the effect that the ordering of natural selection and population regulation in the life cycle has on the resulting population dynamics under the Wright-Fisher model, especially for the evolution of one- and two-locus systems. With weak natural selection, the details of how to order natural selection and population regulation in the life cycle do not matter in the Wright-Fisher model and its diffusion approximation. By contrast, we show that when there is strong natural selection and the population is in linkage disequilibrium, there can be appreciable differences in the resulting population dynamics under the Wright-Fisher model, depending on whether natural selection occurs before or after population regulation in the life cycle. We argue that this effect may be of significance in natural populations subject to gene migration and local selection.

F.Y. supported in part by EPSRC Grant EP/I028498/1.

## 1. Introduction

One of the most important applications of population genetics theory is the inference of genetic and demographic properties of populations. In principle, quantities such as selection coefficients, mutation rates and recombination rates can all be estimated from the gamete frequency data collected at one or multiple time points, at least as products with the effective population size. Accurate inference on the temporal gamete frequency data requires a sound underlying theoretical null model. To date, the Wright-Fisher model and its extensions have been the most popular models in population genetic inference (*e.g*., Sawyer and Hartl, 1992; Yang and Bielawski, 2000; Bustamante et al., 2001; Mathieson and McVean, 2013; Terhorst et al., 2015).

The Wright-Fisher model, developed by Fisher (1930) and Wright (1931), is concerned with a randomly mating population of fixed size evolving in discrete and nonoverlapping generations at a single locus with two alleles, which can be regarded as a simplified version of the life cycle where the next generation is randomly sampled with replacement from an effectively infinite pool of gametes built from equal contributions of all individuals in the current generation. Burger (2000); Ewens (2004) and Durrett (2008) provided an excellent introduction to the Wright-Fisher model, including the versions incorporated with other basic mechanisms of evolution such as mutation and natural selection.

Most theoretical and applied population genetic research rests explicitly or implicitly on the Wright-Fisher model and its extensions, including Kingman’s coalescent (Kingman, 1982), Ewens’ sampling formula (Ewens, 1972; Lessard, 2007) and Kimura’s work on fixation probabilities (Kimura, 1955). Such widespread application is mainly due to the fact that the Wright-Fisher model captures the essential features of the underlying evolutionary model and supplies a concise framework for describing the dynamics of gamete frequencies, even in complex evolutionary scenarios.

In the present work, we are concerned with a randomly mating diploid population of constant size *N* (*i.e*., a population of 2*N* chromosomes) evolving with discrete and nonoverlapping generations under natural selection. The life cycle here is simplified to move through a loop of random mating, natural selection and population regulation, meiosis, random mating, and so forth, which can be summarised in Figure 1. Also, we assume that the entrance of our life cycle is with the population reduced to *N* adults right after a round of population regulation.

**FIGURE 1.**
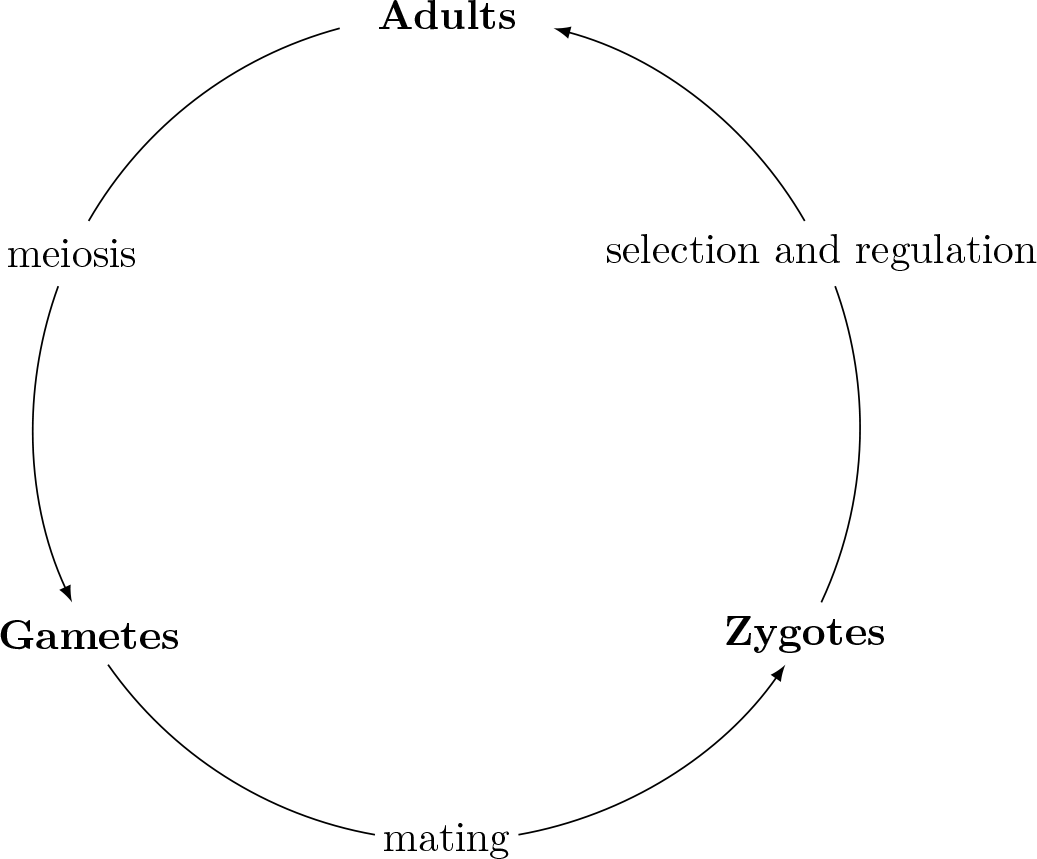
A diagram of the simplified life cycle of diploid populations

Nagylaki (1997) provided different derivations of multinomial-sampling models for random genetic drift at a single locus with multiple alleles in a monoecious or dioecious diploid population for different orders of the evolutionary forces in the life cycle. Prugnolle et al. (2005) found that gene migration occurring before and after asexual reproduction in the life cycle of monoecious trematodes can have different effects on a finite island model, depending on values of the other genetic parameters. A natural question that arises here is whether different sequences of the evolutionary processes in the life cycle can cause different behaviour in the Wright-Fisher model and its extensions.

As shown in Figure 1, while developing into adults, zygotes undergo natural selection, and at the same time, *N* zygotes are randomly sampled with replacement to survive to adulthood by population regulation (*i.e*., genetic drift here acts through population regulation). We are thereby left with two potential choices to order natural selection and population regulation in the life cycle when we set up a Wright-Fisher model, *i.e*., natural selection occurs before or after population regulation, which we refer to as the *ordering* in the following. That is, there are two possible sequences of the evolutionary processes in the life cycle: meiosis, random mating, natural selection, population regulation (*abbr. sr*); and natural selection, meiosis, random mating, population regulation (*abbr. rs*). The objective of the present work is mainly to address the question of whether different *orderings* have different effects on the behaviour of the Wright-Fisher model.

The remainder of this work is organised as follows. In Section 1, we set up a generic version of the Wright-Fisher model for the life cycle illustrated in Figure 1, and provide detailed derivations for the evolution of one- and two-locus systems under natural selection, respectively. In Section 3, we investigate the effect of the *ordering* on the behaviour of the Wright-Fisher model with the aid of Monte Carlo simulations. Finally, in Section 4, we conclude with some discussion on how the *ordering* affects the behaviour of the diffusion approximation of the Wright-Fisher model and offer some further perspectives.

## 2. Wright-Fisher model

In this section, we give a derivation of a generic version of the Wright-Fisher model for the life cycle illustrated in Figure 1, then provide detailed formulations for the evolution of one- and two-locus systems under natural selection for the two *orderings*, *sr* and *rs*.

### 2.1. Wright-Fisher Model with Selection

Consider a monoecious population of *N* randomly mating diploid individuals evolving with discrete and nonoverlapping generations under natural selection. We assume that there are two alleles segregating at each autosomal locus, and the population size is fixed. Let 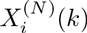 be the frequency of gamete *i* in *N* adults of generation *k* ∈ ℕ and ***X**^(*N*)^(*k*)* denote the vector with frequencies of all possible gametes. We can study the population dynamics under natural selection in terms of the changes in gamete frequencies from generation to generation.

To determine the transition of gamete frequencies from one generation to the next, however, we need to investigate how the mechanisms of evolution (*e.g*., natural selection) alter the genotype frequencies at intermediate stages of the life cycle. Let 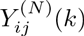 be the frequency of the (ordered) genotype made up of gametes *i* and *j* in *N* adults of generation *k* and ***Y***^(*N*)^(*k*) designate the vector with frequencies of all possible genotypes. Under the assumption of random mating, the genotype frequency is equal to the product of the corresponding gamete frequencies,

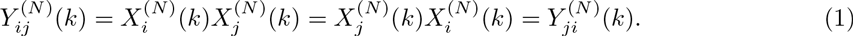

As illustrated in Figure 1, the life cycle moves through a loop of random mating, natural selection and population regulation, meiosis, random mating, and so forth. That is, there are three potential mechanisms of evolutionary change, genetic recombination (during meiosis), natural selection and population regulation, in the life cycle required to be accounted for here. From the intuition that comes from the Wright-Fisher model, we assume that natural selection and genetic recombination occur in an effectively infinite population so can be treated deterministically (see Hamilton, 2011). Furthermore, suppose that population regulation (*i.e*., genetic drift) acts in a similar manner to the Wright-Fisher sampling introduced by Fisher (1930) and Wright (1931). In other words, population regulation corresponds to randomly drawing *N* zygotes with replacement from an effectively infinite population to become new adults in the next generation, consequently completing the life cycle. Therefore, given the genotype frequencies ***Y***^(*N*)^(*k*) = *y*, the genotype frequencies in the next generation satisfy

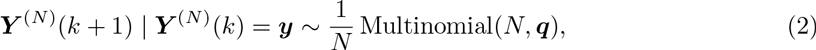

where ***q*** is the vector with frequencies of all possible genotypes of an effectively infinite population after a single generation of the possible evolutionary forces (except genetic drift) at intermediate stages of the life cycle such as natural selection and genetic recombination. The explicit expression of the sampling probabilities ***q*** will be given in the following two sections for the evolution of one- and two-locus systems under natural selection, respectively.

To ease notations, we introduce a function 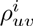 of three variables *u*, *v* and *i*, defined as

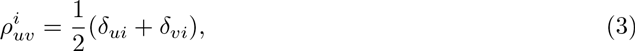

where *δ_ui_* and *δ_vi_* denote the Kronecker delta functions. We can then express the frequency of gamete *i* in terms of the corresponding genotype frequencies as

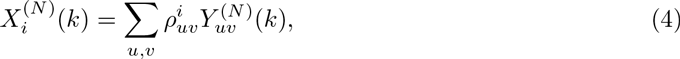

and the transition probabilities of the gamete frequencies from one generation to the next can be easily obtained from Eqs. (2)) and (4). Here we refer to the process of gamete frequencies ***X***^(*N*)^ = {***X***^(*N*)^(*k*): *k* ∈ ℕ} as the Wright-Fisher model with selection.

### 2.2. one-locus Wright-Fisher Model with Selection

Now we consider a monoecious population of *N* randomly mating diploid individuals at a single autosomal locus 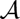, evolving under natural selection in discrete and nonoverlapping generations. Suppose that there are two possible allele types at locus 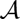, labelled 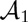 and 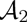, respectively. As mentioned in Section 2.1, it remains to formulate the sampling probabilities ***q*** for the one-locus Wright-Fisher model with selection, which is the vector with frequencies of all possible genotypes of an effectively infinite population after a single generation of the potential evolutionary forces (except genetic drift) at intermediate stages of the life cycle. In the one-locus model, due to the absence of genetic recombination during meiosis, the two *orderings*, *sr* and *rs*, can be reduced to one identical order of the evolutionary forces that alter the sampling probabilities ***q***. That is, ***q*** is the vector with frequencies of all possible genotypes of an effectively infinite population after a single generation of natural selection.

With a single diallelic locus, there are four possible (ordered) genotypes that result from random union of two alleles. We assume that natural selection acts only through viability differences and denote the fitness of the genotype formed by alleles *i* and *j* by *w_ij_* (*i.e*., the probability that zygotes of the corresponding genotype survive to adulthood). According to Hamilton (2011), we can formulate the frequency of the genotype formed by alleles *i* and *j* of an effectively infinite population after a single generation of natural selection as

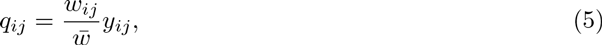

where

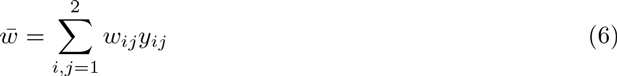

is the mean fitness.

From Eqs. (5)-(6), we see that the transition probabilities of the gamete frequencies from one generation to the next only depend on the gamete frequencies in the current generation. Combining with Eqs. (2) and (4), we find that the process of gamete frequencies ***X***^(*N*)^ is a time-homogeneous Markov process evolving in the state space

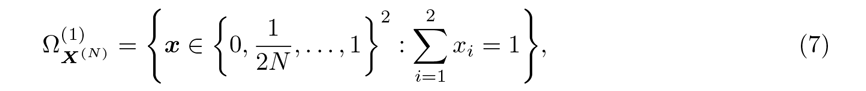

which we call the one-locus Wright-Fisher model with selection.

### 2.3. Two-locus Wright-Fisher Model with Selection

Consider a monoecious population of *N* randomly mating diploid individuals at two linked autosomal loci named 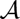 and 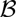, each segregating into two alleles, 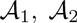 and 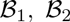, evolving under natural selection with discrete and nonoverlapping generations. There are four possible types of gametes 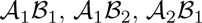 and 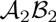 called gametes 1, 2, 3 and 4, respectively. As we have stated in Section 2.1, we need to formulate the sampling probabilities ***q*** for the two-locus Wright-Fisher model with selection, which is the vector with frequencies of all possible genotypes of an effectively infinite population after a single generation of the potential evolutionary forces (except genetic drift) at intermediate stages of the life cycle. In the two-locus model, due to the existence of genetic recombination during meiosis, we can reduce the two *orderings*, *sr* and *rs*, to two different orders of the evolutionary forces that alter the sampling probabilities, denoted by ***q***^(*sr*)^ and ***q***^(*rs*)^, respectively, where ***q***^(*sr*)^ denotes the vector with frequencies of all possible genotypes of an effectively infinite population after a single generation of genetic recombination and natural selection when natural selection occurs before population regulation, and ***q***^(*rs*)^ denotes the vector with frequencies of all possible genotypes of an effectively infinite population after a single generation of genetic recombination and natural selection when natural selection occurs after population regulation.

With two diallelic loci, there are sixteen possible (ordered) genotypes that result from random union of four gametes. We assume that natural selection acts only through viability differences, and denote the fitness of the genotype formed by gametes *i* and *j* by *w_ij_*. We designate the recombination rate between the two loci by *r*, which is in the range 0 ≤ *r* ≤ 0.5. To ease notations, we introduce a vector of auxiliary variables ***η*** = (*η*_1_,*η*_2_, *η*_3_, *η*_4_), where *η*_1_ = *η*_4_ = −1 and *η*_2_ = *η*_3_ = 1. From Hamilton (2011), the frequency of the genotype formed by gametes *i* and *j* of an effectively infinite population after a single generation of natural selection only is

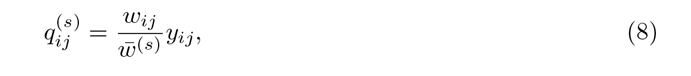

where

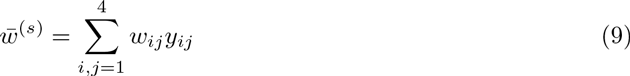

is the mean fitness, and the frequency of the genotype formed by gametes *i* and *j* of an effectively infinite population after a single generation of genetic recombination only is

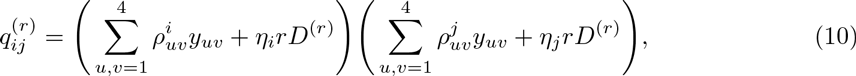

where

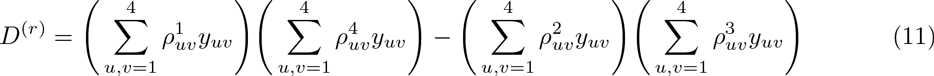

is the standard measure of linkage disequilibrium introduced by Lewontin and Kojima (1960), describing the level of the non-random association of alleles at two loci here.

With the *ordering sr*, we calculate the genotype frequencies of an effectively infinite population after a single generation of genetic recombination using Eqs. (10) and (11) first then incorporate natural selection by replacing *y_ij_* with 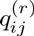 in Eqs. (8) and (9). We can then express the frequency of the genotype formed by gametes *i* and *j* of an effectively infinite population after a single generation of genetic recombination and natural selection when natural selection occurs before population regulation as

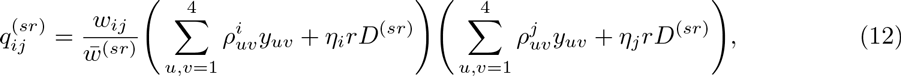

where

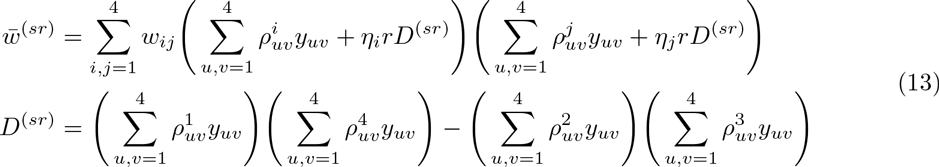

are the corresponding mean fitness and the standard measure of linkage disequilibrium. We refer to the Wright-Fisher model with the *ordering sr* as the *sr*-Wright-Fisher model.

With the *ordering rs*, we calculate the genotype frequencies of an effectively infinite population after a single generation of natural selection using Eqs. (8) and (9) first then incorporate genetic recombination by replacing *y_ij_* with 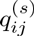 in Eqs. (10) and (11). We can then express the frequency of the genotype formed by gametes *i* and *j* of an effectively infinite population after a single generation of genetic recombination and natural selection when natural selection occurs after population regulation as

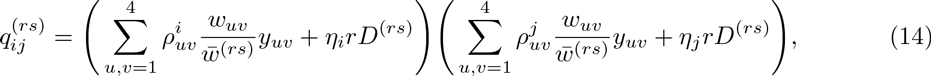

where

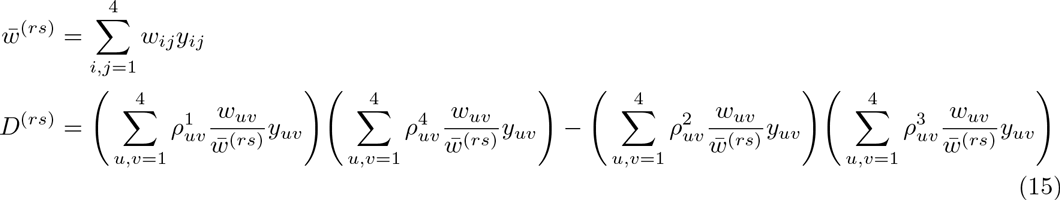

are the corresponding mean fitness and the standard measure of linkage disequilibrium. We refer to the Wright-Fisher model with the *ordering rs* as the *rs*-Wright-Fisher model.

From Eqs. (12)-(13) and (14)-(15), we see that the transition probabilities of the gamete frequencies from one generation to the next depend only on the gamete frequencies in the present generation in both of the two *orderings, sr* and *rs*, but take on different forms. Combining with Eqs. (2) and (4), we find that the process of gamete frequencies ***X***^(*N*)^ is a time-homogeneous Markov process evolving in the state space

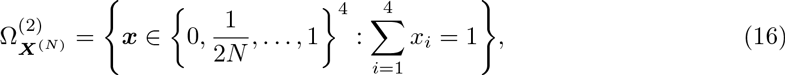

which we call the two-locus Wright-Fisher model with selection.

## 3. Effects of the Ordering of Selection and Regulation

In this section, we are concerned with the question of whether different *orderings* have different effects on the behaviour of the Wright-Fisher model, especially for the evolution of one- and two-locus systems under natural selection. Since the Wright-Fisher model can be completely determined by the probability of the transition, our problem can reduce to investigating the effect of the *ordering* on the behaviour of the Wright-Fisher model in terms of transition probabilities. As demonstrated in Section 2, the transition probabilities of a single locus are unchanged, but correlation between loci can be affected. We therefore only consider the population dynamics under natural selection at two linked loci here. Notice that we illustrate how the *ordering* affects the behaviour of the Wright-Fisher model with the 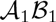 gamete in detail in the following and expect other gametes to behave in a similar manner.

Before exploring the effect of the *ordering* on the behaviour of the Wright-Fisher model, we need make further assumptions on the mechanism of natural selection. From Gillespie (2010), we employ the relative fitness rather than the absolute fitness, and a common notational convention of the relative fitness for diploid populations at a single locus can be defined as follows: genotypes 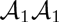, 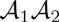 and 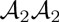 at a given locus 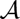 have relative fitnesses 1, 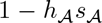 and 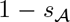, respectively, where *s* is the selection coefficient and *h* is the dominance parameter. We consider the simple case of directional selection with 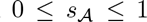, which implies that the 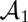 allele is the type favoured by natural selection. Suppose that the dominance parameter is in the range 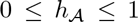, *i.e*., general dominance. Furthermore, we assume that relative fitnesses of the two-locus genotypes are determined multiplicatively from relative fitnesses at individual loci, *e.g*., the relative fitness of the 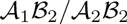 genotype is 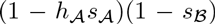, which means that there is no position effect, *i.e*., coupling and repulsion double heterozygotes have the same relative fitness, *w*_14_ = *w*_23_ = 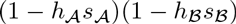.

Let *π*_1_(*k*, *x*) denote the vector with probabilities of the transition of gamete 1 (*i.e*., the 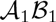 gamete) from the initial state of the population *x* over *k* generations, where *x* is the vector with initial frequencies of four possible gametes. We apply the Hellinger distance between the two transition probabilities 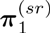 and 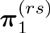 to quantify the difference in the behaviour of the *sr*- and *rs*-Wright-Fisher models and consider the dynamics of the Hellinger distance over time with varying genetic parameters. From Le Cam and Yang (2000), theHellinger distancebetweenthe two transition probabilities 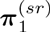 and 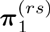 can be defined by the quantity

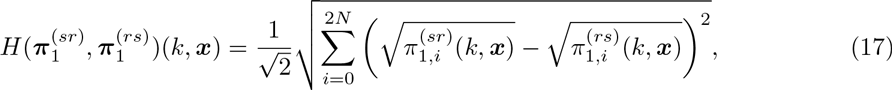

where *π*_1*i*_(*k*, *x*) is the probability of the transition of the frequency of the 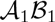 gamete to *i*/(2*N*) from the population of the initial state *x* over *k* generations.

### 3.1. Monte Carlo Simulation Studies

Given the difficulties in analytically computing the probability of the transition, especially for the population evolving over a long time period, we resort to Monte Carlo simulations here, which enable us to get an empirical transition probabilities 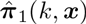, defined by

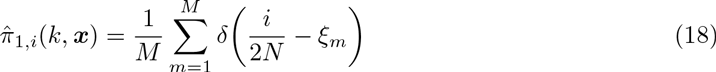

for *i* = 0,1,…, 2*N*, where *ξ_m_* is the *m*-th realisation of the frequency of the 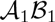 gamete simulated under the Wright-Fisher model from the population of the initial state *x* over *k* generations and *M* is the total number of realisations in the sample. Combing with Eqs. (17) and (18), we can evaluate the Monte Carlo approximation of the Hellinger distance 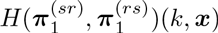 as

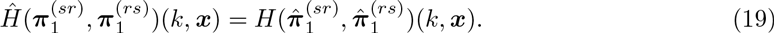

In the following illustration, we investigate the effect of the *ordering* on the behaviour of the Wright-Fisher model from the dynamics of the Hellinger distance 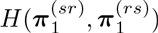 over time with varying genetic parameters according to the population size: large populations and small populations, and the dynamics of the Hellinger distance 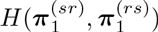 is simulated through Monte Carlo. Due to symmetry of the two loci, we fix the selection coefficient 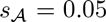 and dominance parameter 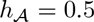 at locus 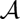 in the following.

#### 3.1.1. *Large Populations.*

We are concerned with a large population evolving under natural selection at two linked loci according to the Wright-Fisher model demonstrated in Section 3, in which we take the population size to be *N* = 5000. To investigate how the *ordering* affects the behaviour of the Wright-Fisher model, we simulate the dynamics of the Hellinger distance 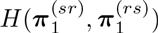 over time with varying genetic parameters, where we adopt the number of realisations *M* = 10^5^. We present the results for the following three particular scenarios according to the types of gene action at locus 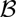, respectively: 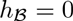 (completely dominant), 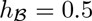 (completely additive) and 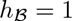 (completely recessive).

In Figures 2–4, we show the dynamics of the Hellinger distance 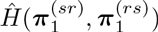 over 400 generations when gene action at locus 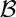 is completely dominant, additive and recessive, respectively. From the middle columns of Figures 2–4, the Hellinger distance 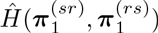 evolving over time is always close to 0 when the population is initially in linkage equilibrium. However, the left and right columns of Figures 2–4 illustrate that when the population is initially far away from linkage equilibrium, the Hellinger distance 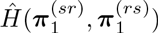 evolving over time is no longer close to 0, especially for large selection coefficients and recombination rates. Intuitively, the effect of the *ordering* on the behaviour of the two-locus Wright-Fisher model does exist in large populations.

**FIGURE 2.**
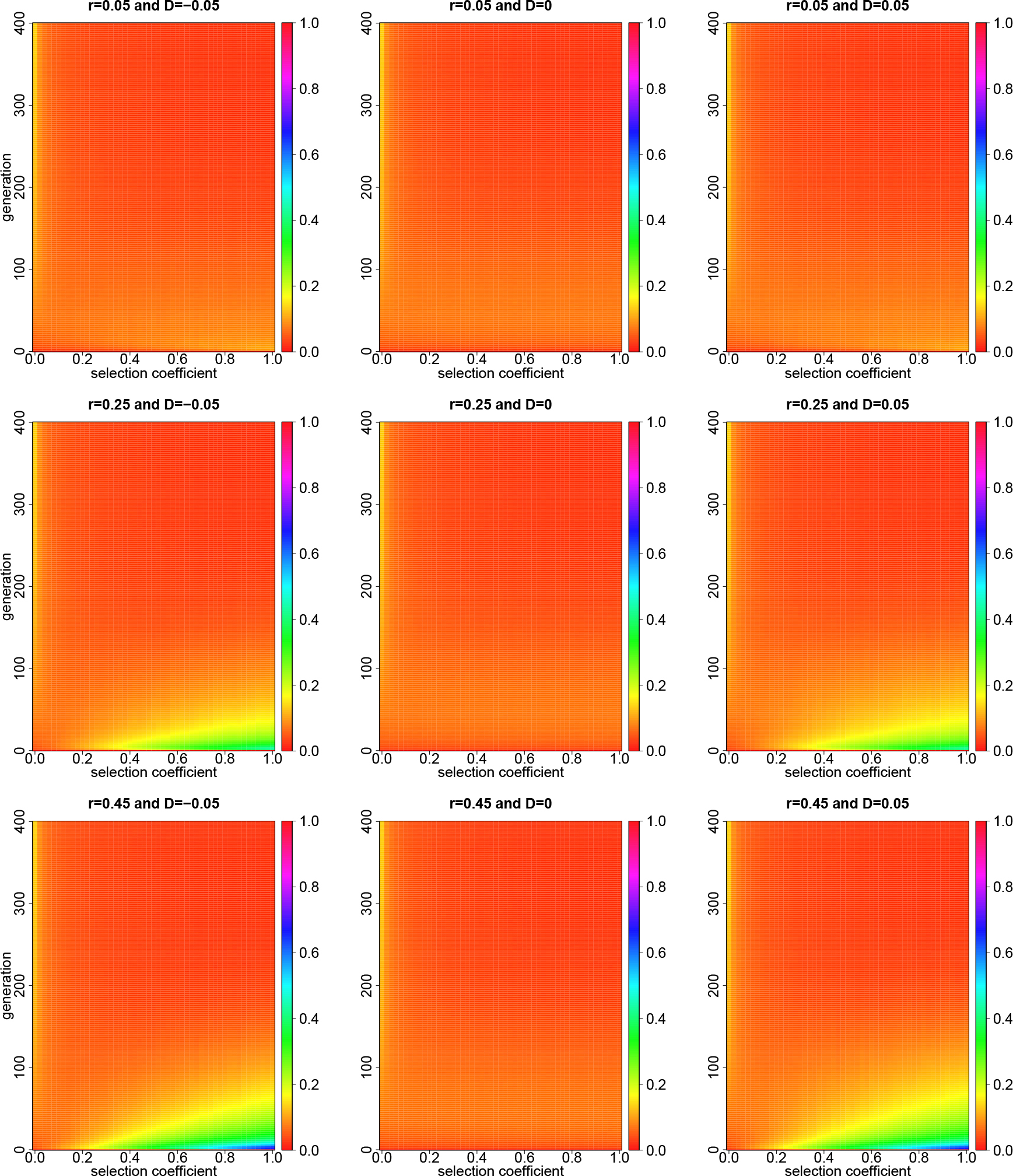
Dynamics of the Hellinger distance 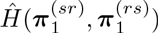 with varying selection coefficients 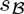 over 400 generations for a large population when the gene action at locus 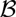 is completely dominant. The Hellinger distance 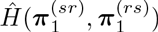 is evaluated from Monte Carlo simulations with 10^5^ independent realisations. The other parameters are *N* = 5000, 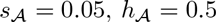 and 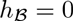.

**FIGURE 3.**
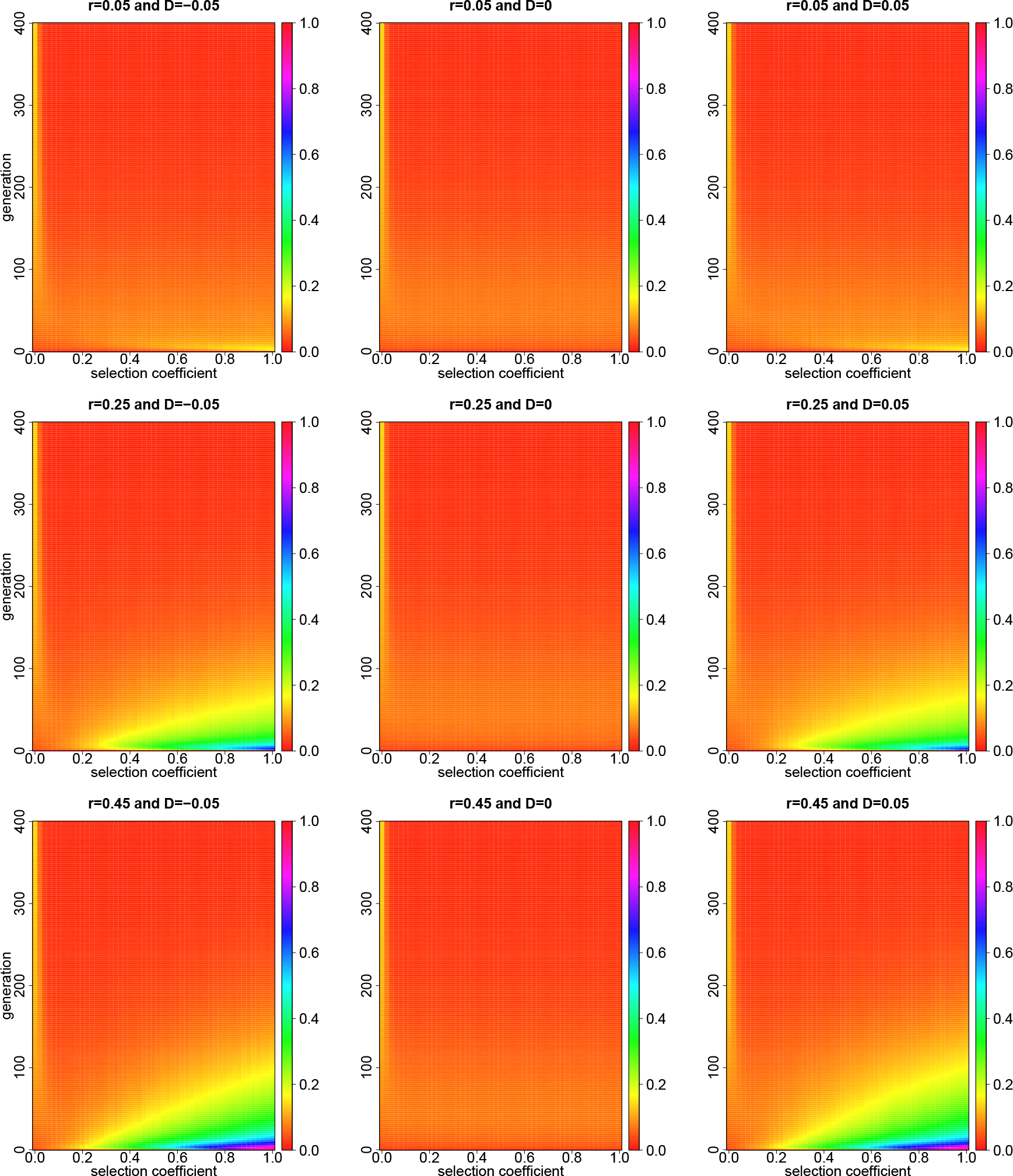
Dynamics of the Hellinger distance 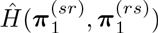 with varying selection coefficients 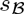 over 400 generations for a large population when the gene action at locus 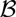 is completely additive. The Hellinger distance 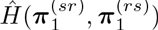 is evaluated from Monte Carlo simulations with 10^5^ independent realisations. The other parameters are *N* = 5000, 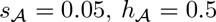 and 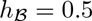

**FIGURE 4.**
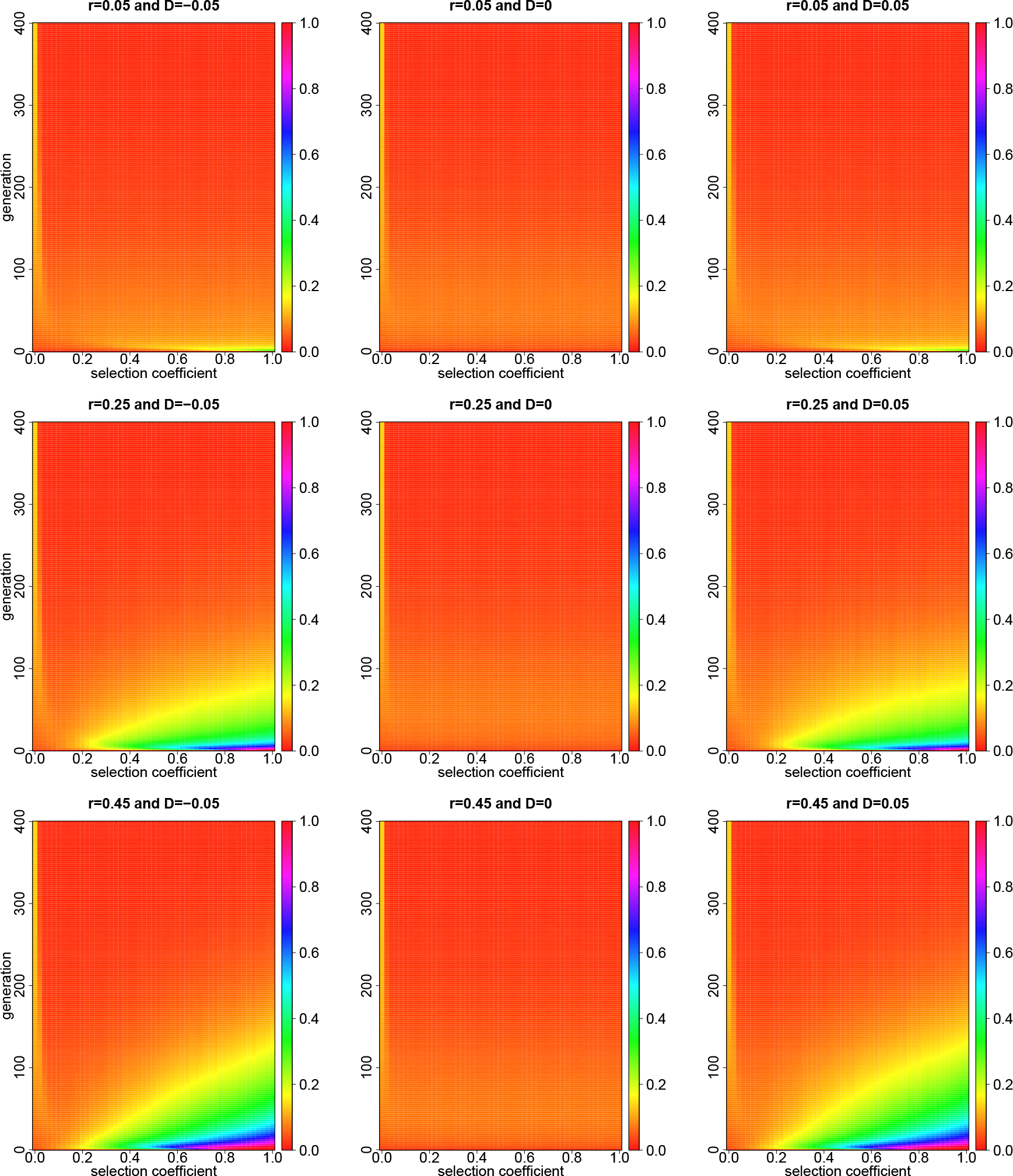
Dynamics of the Hellinger distance 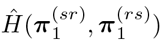 with varying selection coefficients 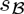 over 400 generations for a large population when the gene action at locus 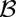 is completely recessive. The Hellinger distance 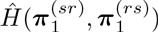 is evaluated from Monte Carlo simulations with 10^5^ independent realisations. The other parameters are *N* = 5000, 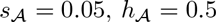 and 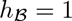.

Comparing the middle columns of Figures 2–4 with their left and right columns, we find that whether the *ordering* affects the behaviour of the Wright-Fisher model depends largely on the linkage disequilibrium. We therefore consider the dynamics of linkage disequilibrium over time before investigating how the *ordering* affects the behaviour of the Wright-Fisher model. In Figure 5, we simulate the dynamics of the transition probabilities of linkage disequilibrium over time with given genetic parameters. As illustrated in Figure 5, the probability of the transition of linkage disequilibrium becomes concentrated on a narrower and narrower range of possible values centred around 0 from generation to generation until completely fixed at 0. As we have stated in Section 2.1, there are three potential mechanisms of evolutionary change, genetic recombination, natural selection and population regulation (genetic drift), in the life cycle, which may affect the dynamics of linkage disequilibrium over time. Genetic recombination always works towards linkage equilibrium due to genetic recombination generating new gamete types to break down non-random genetic associations (Ridley, 2004). Natural selection alone can not move the population far away from linkage equilibrium under the assumption that relative fitnesses are multiplicative (Slatkin, 2008). Genetic drift can destroy linkage equilibrium and create many genetic associations since genetic drift leads to the random change in gamete frequencies (Ridley, 2004). However, in large populations, the amount of the change in gamete frequencies caused by genetic drift is quite small. So once linkage equilibrium has been reached, the large population will not tend to be far away from linkage equilibrium.

**FIGURE 5.**
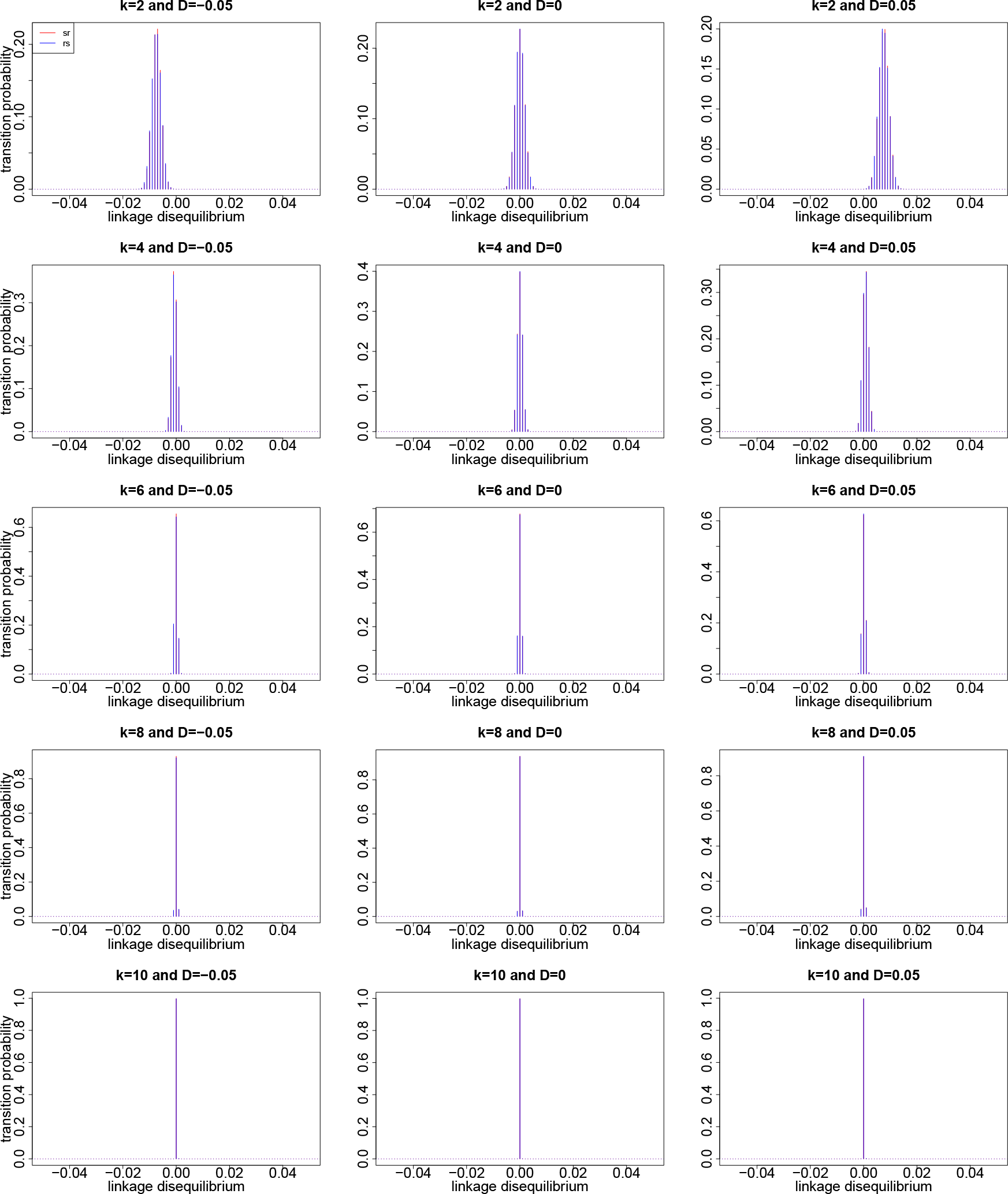
Comparison of the empirical transition probabilities of linkage disequilibrium for the two *orderings sr* and *rs* in generations *k* = 2, 4, 6, 8,10 for a large population. The empirical transition probabilities are evaluated from Monte Carlo simulations with 10^5^ independent realisations. In this illustration, we take *N* = 5000, 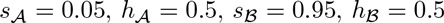 and *r* = 0.45.

From Eqs. (12)-(13) and (14)-(15), provided that the population evolving over time is always close to linkage equilibrium, the amount of the change in gamete frequencies caused by genetic recombination is negligible. The two-locus Wright-Fisher model for each locus segregating two alleles thereby becomes similar to the one-locus Wright-Fisher model for a single locus segregating four alleles, where each gamete is analogous to a single allele. Therefore, once linkage equilibrium has been reached, the effect of the *ordering* on the behaviour of the two-locus Wright-Fisher model is negligible in large populations.

Now we discuss how the *ordering* affects the behaviour of the two-locus Wright-Fisher model. As mentioned above, when the population is initially in linkage equilibrium, the effect of the *ordering* on the behaviour of the Wright-Fisher model is negligible in large populations, which is confirmed with the middle columns of Figures 2–4. However, when the population is initially far away from linkage equilibrium, the effect of the *ordering* on the behaviour of the Wright-Fisher model does exist in large populations and lasts until linkage equilibrium has been reached, which is no longer negligible, as shown in the left and right columns of Figures 2–4.

In Figure 6, we illustrate the dynamics of the transition probabilities 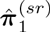 and 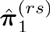 over time with given genetic parameters for the population initially in negative linkage disequilibrium, linkage equilibrium and positive linkage disequilibrium, respectively, and provide the corresponding dynamics of the transition probabilities 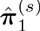 over time as a reference, where 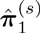 designates the empirical probability of the transition of the 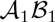 gamete under natural selection at two completely linked loci for the Wright-Fisher model, *i.e*., no recombinants are produced. As shown in Figure 6, it is clear that the *sr*- and *rs*-Wright-Fisher models have different rates of the change in the frequency of the 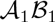 gamete approaching fixation when the population is initially far away from linkage equilibrium, which leads to the difference in the behaviour of the Wright-Fisher model between two different *orderings*, as illustrated in the left and right columns of Figures 2–4.

**FIGURE 6.**
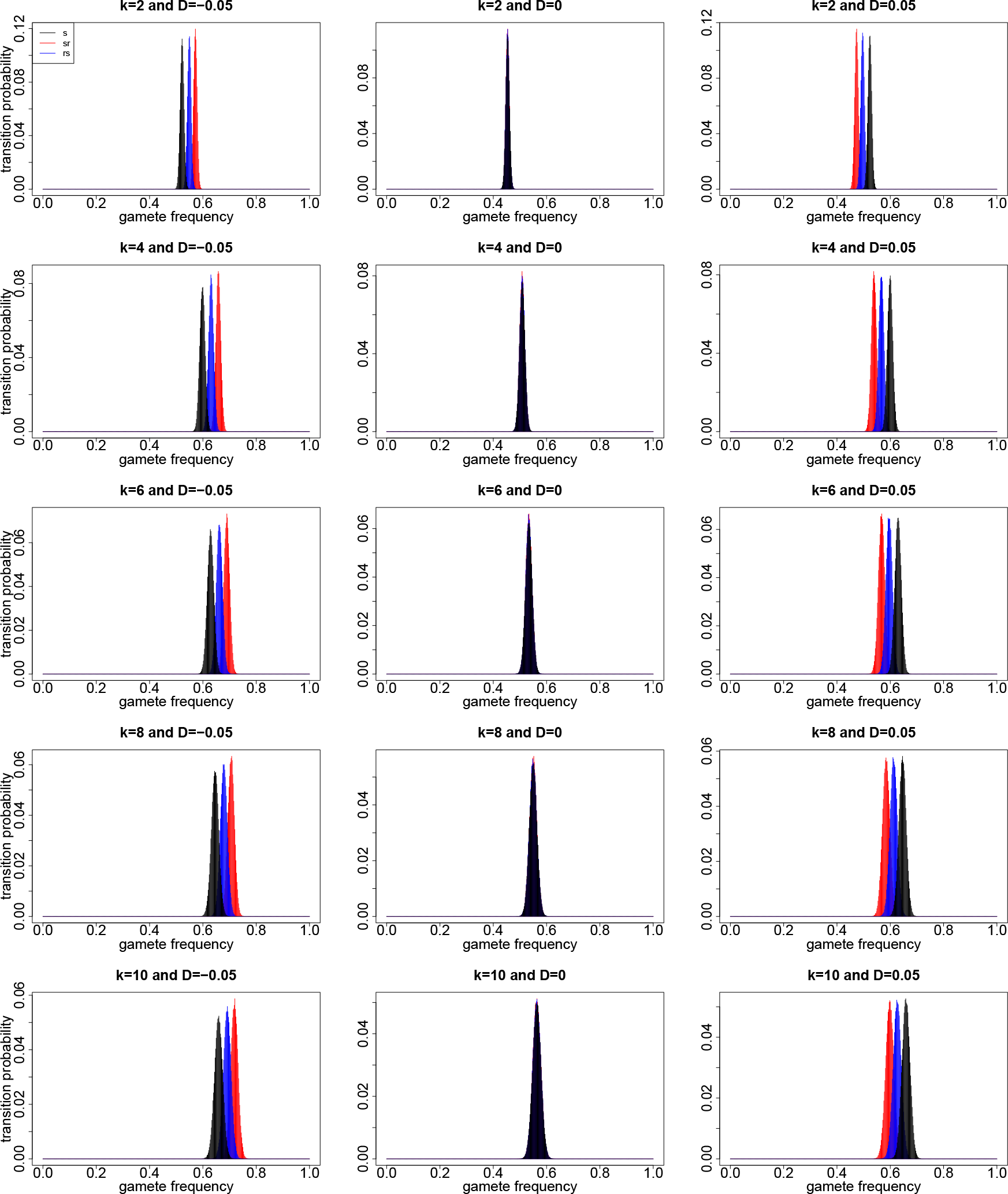
Comparison of the empirical transition probabilities of the 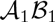 gamete for the two *orderings sr* and *rs* in generations *k* = 2, 4, 6, 8, 10 for a large population. The empirical transition probabilities are evaluated from Monte Carlo simulations with 10^5^ independent realisations. In this illustration, we take *N* = 5000, 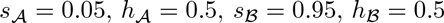 and *r* = 0.45.

More specifically, as shown in the left column of Figure 6, when the population is in negative linkage disequilibrium, the Wright-Fisher model of two linked loci drives a more rapid approaching to the fixation of the 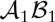 gamete than the Wright-Fisher model of two completely linked loci, which is due to the fact that genetic recombination reinforces the change in gamete frequencies caused by natural selection for negative linkage disequilibrium (see Hamilton, 2011). Furthermore, the *sr*-Wright-Fisher model promotes a more rapid approaching to the fixation of the 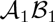 gamete than the *rs*-Wright-Fisher model when the population is in negative linkage disequilibrium, which implies that genetic recombination affecting natural selection at two linked loci is more significant in the *sr*-Wright-Fisher model than that in the *rs*-Wright-Fisher model for negative linkage disequilibrium. These results are also confirmed by the right column of Figure 6 for the population in positive linkage disequilibrium. Therefore, in large populations, the difference in the behaviour of the two-locus Wright-Fisher model between two different *orderings* results from the different performance of genetic recombination affecting natural selection for different *orderings* in the Wright-Fisher model, and genetic recombination has a more significant effect on natural selection at two linked loci when natural selection occurs before population regulation. Moreover, from Figures 2–4, we find that this effect exponentially increases as genetic recombination increases.

#### 3.1.2. *Small Populations.*

Now we consider a small population evolving under natural selection at two linked loci according to the Wright-Fisher model demonstrated in Section 3. We take the population size to be *N* = 50. To investigate how the *ordering* affects the behaviour of the Wright-Fisher model, we simulate the dynamics of the Hellinger distance 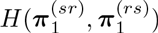 over time with varying genetic parameters, where we adopt the number of realisations *M* = 10^5^. Similarly, we present the results for completely dominant, additive and recessive gene action at locus 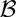, respectively.

In Figures 7-9, we illustrate the dynamics of the Hellinger distance 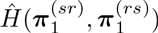 over 400 generations when gene action at locus 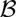 is completely dominant, additive and recessive, respectively. Figures 7-9 confirm the results we have obtained for large populations, but the Hellinger distance 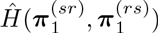 in small populations is much smaller than that in large populations. That is, the effect of the *ordering* on the behaviour of the Wright-Fisher model is significantly weakened in small populations.

**FIGURE 7.**
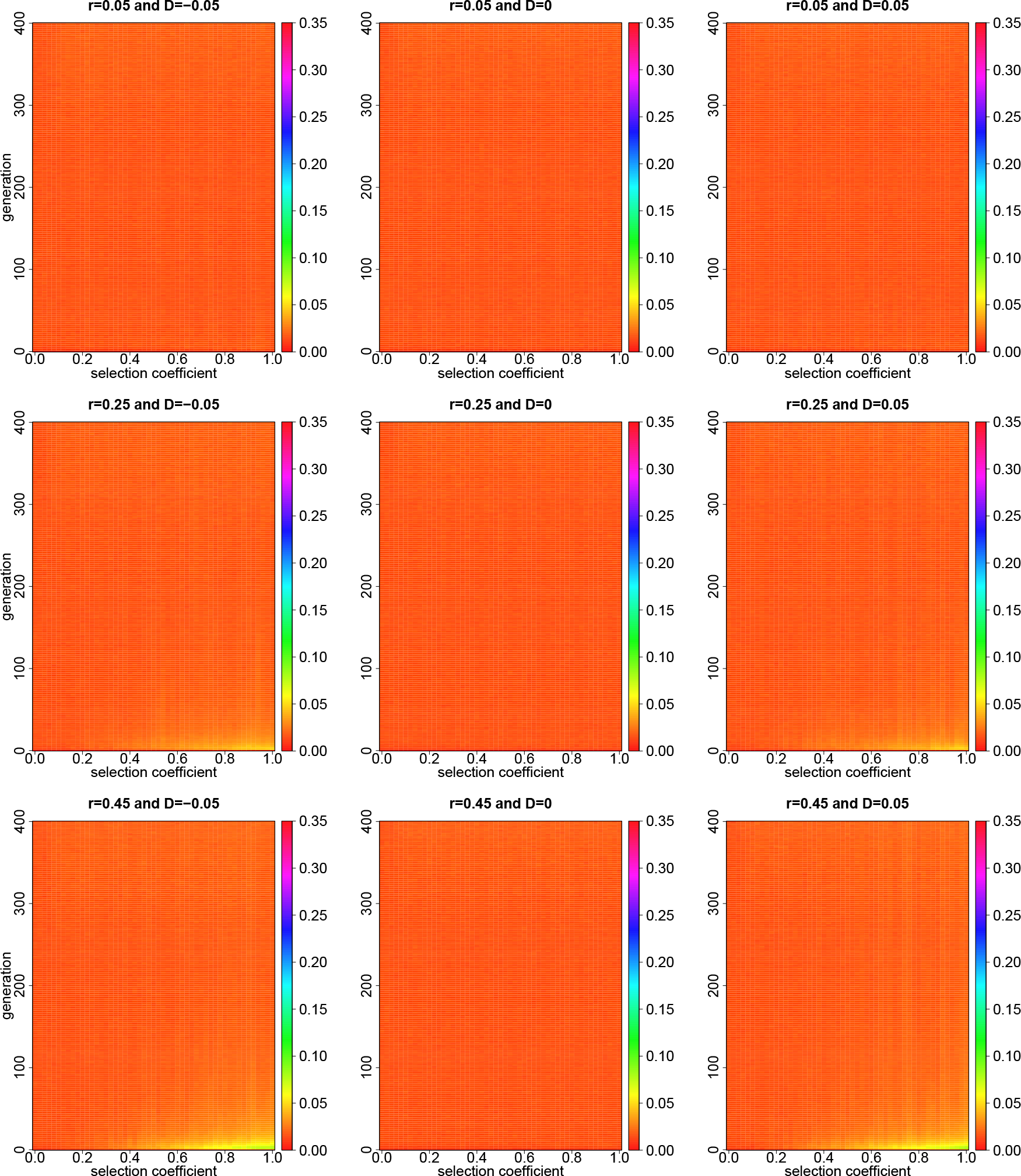
Dynamics of the Hellinger distance 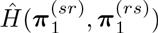 with varying selection coefficients 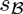 over 400 generations for a small population when the gene action at locus 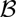 is completely dominant. The Hellinger distance 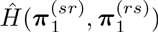 is evaluated from Monte Carlo simulations with 10^5^ independent realisations. The other parameters are *N* = 50,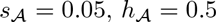 and 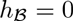.

**FIGURE 8.**
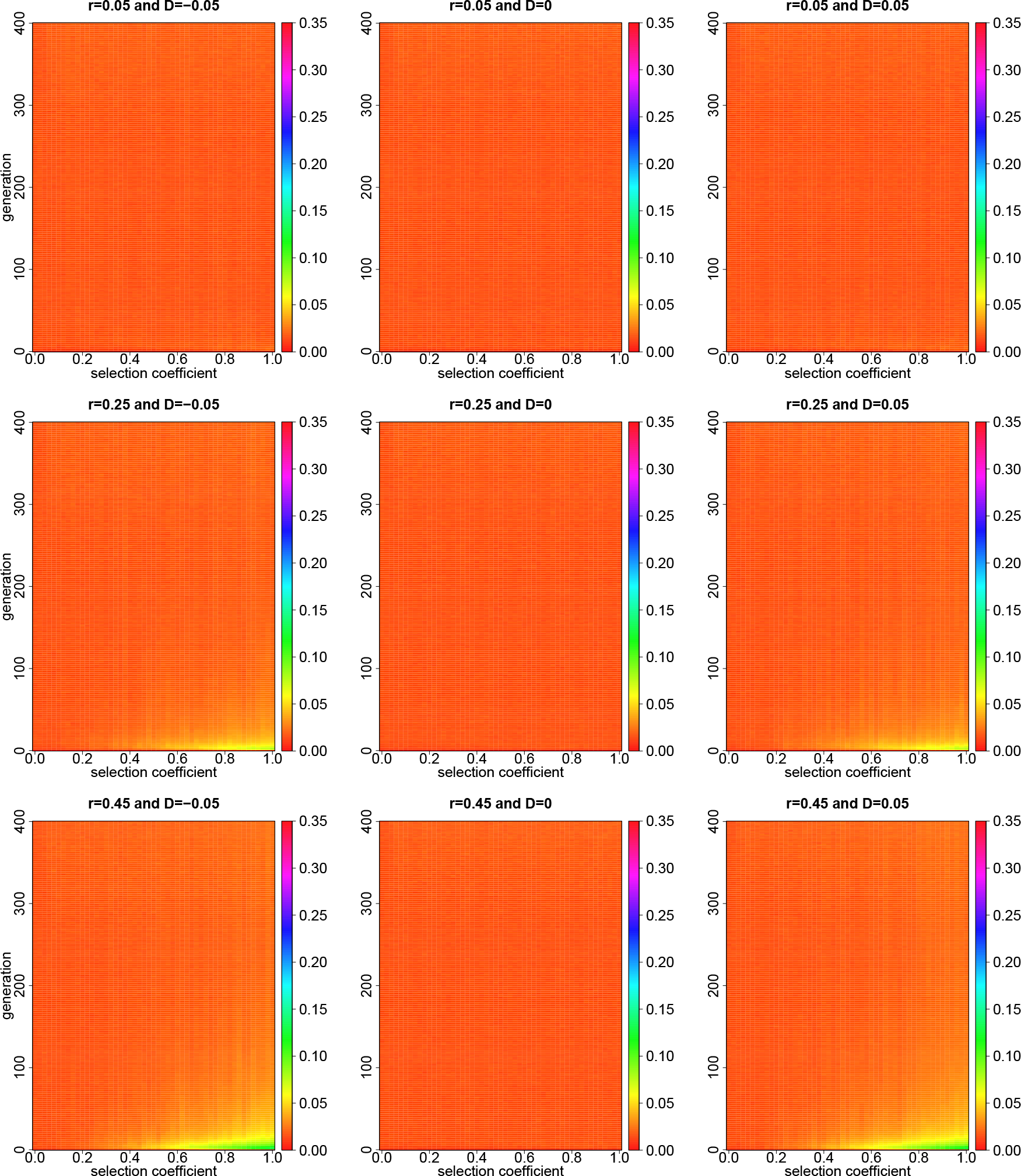
Figure 8. Dynamics of the Hellinger distance 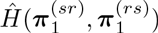 with varying selection coefficients 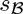 over 400 generations for a small population when the gene action at locus 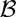 is completely additive. The Hellinger distance 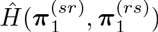 is evaluated from Monte Carlo simulations with 10^5^ independent realisations. The other parameters are *N* = 50, 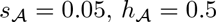 and 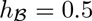.

**FIGURE 9.**
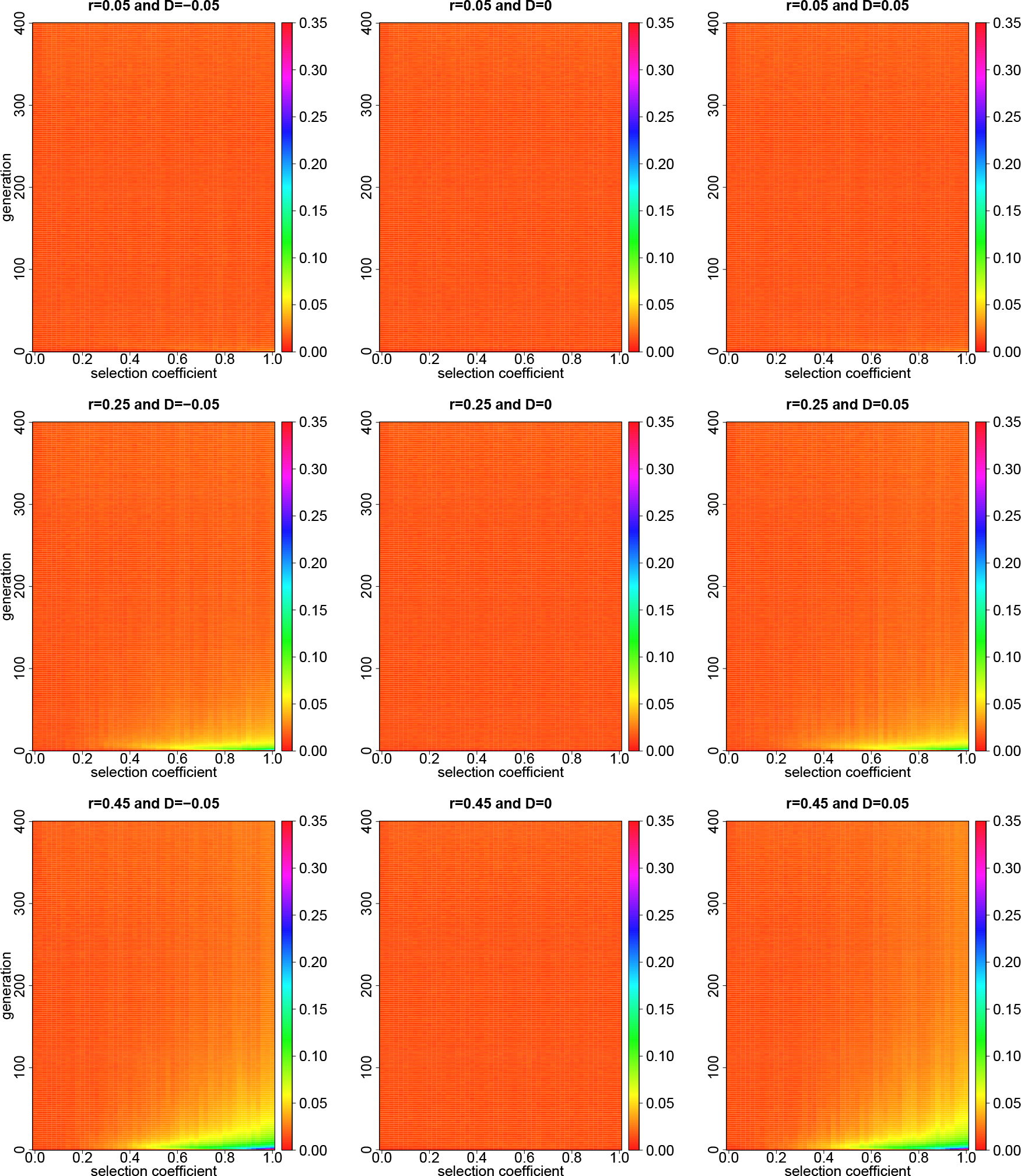
Dynamics of the Hellinger distance 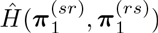,with varying selection coefficients 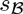 over 400 generations for a small population when the gene action at locus 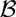 is completely recessive. The Hellinger distance 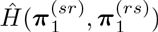 is evaluated from Monte Carlo simulations with 10^5^ independent realisations. The other parameters are *N* = 50, 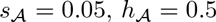 and 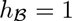.

**FIGURE 10.**
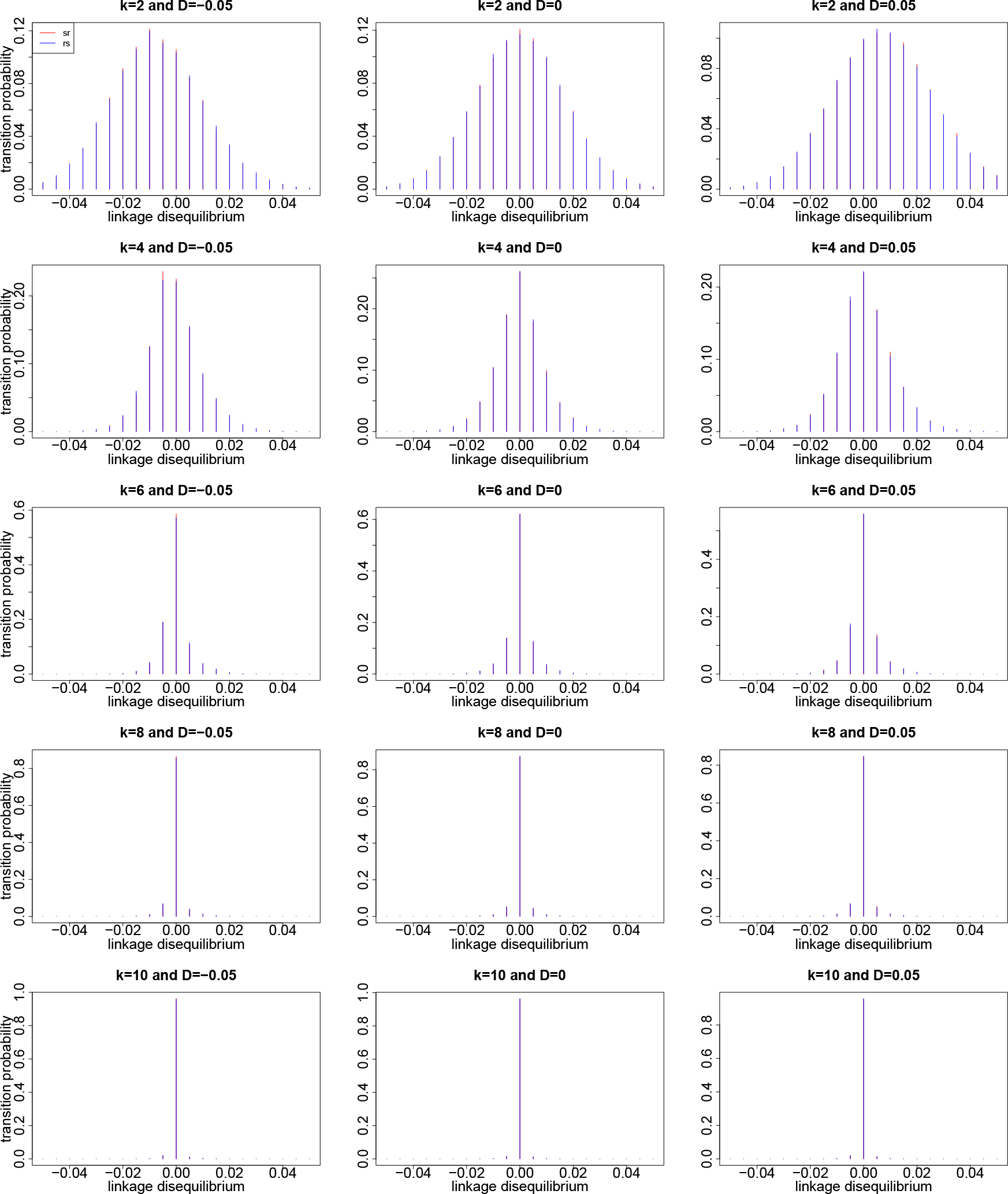
Comparison of the empirical transition probabilities of linkage disequilibrium for the two *orderings sr* and *rs* in generations *k* = 2, 4, 6, 8,10 for a small population. The empirical transition probabilities are evaluated from Monte Carlo simulations with 10^5^ independent realisations. In this illustration, we take *N* = 50, 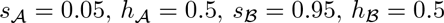 and *r* = 0.45.

Similarly, we simulate the dynamics of the transition probabilities of linkage disequilibrium over time in Figure 10 and the dynamics of the transition probabilities 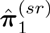 and 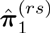 over time in Figure 11 for small populations. Comparing Figures 10-11 with 5-6, we find that the breadth of the transition probabilities widens with decreasing population size due to the strong effect of genetic drift in small populations. That is, the stronger genetic drift in small populations leads to an increase in sampling error, which reduces the effect of the *ordering* on the behaviour of the Wright-Fisher model. This stronger genetic drift in small populations also results in persistent linkage disequilibrium, and the population requires more generations to reach linkage equilibrium (see Figure 10). Hence, the effect of the *ordering* on the behaviour of the Wright-Fisher model lasts longer in small populations than that in large populations.

**FIGURE 11.**
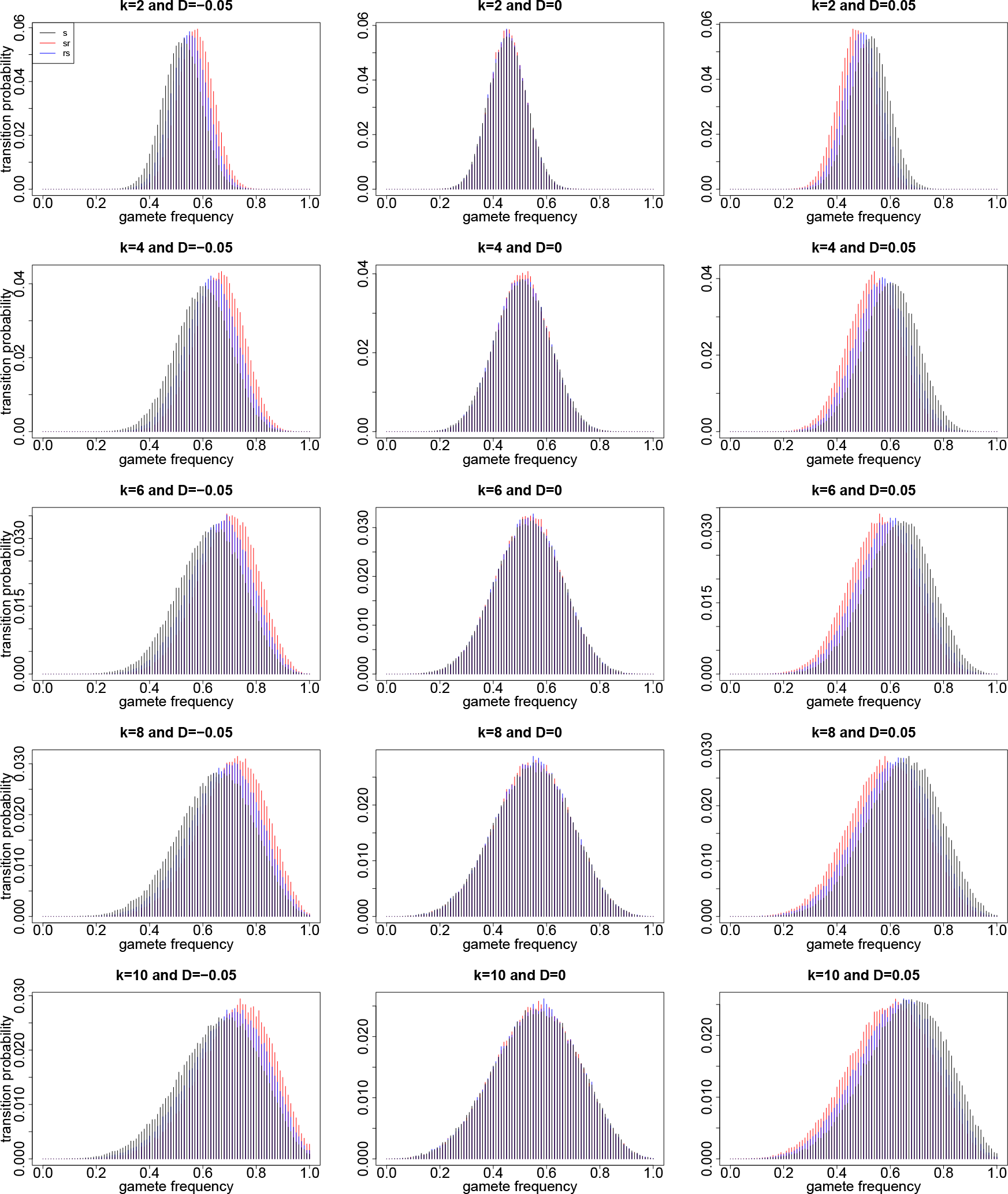
Comparison of the empirical transition probabilities of the 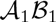 gamete for the two *orderings sr* and *rs* in generations *k* = 2,4, 6, 8,10 for a small population. The empirical transition probabilities are evaluated from Monte Carlo simulations with 10^5^ independent realisations. In this illustration, we take *N* = 50, 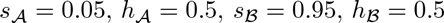 and *r* = 0.45.

In conclusion, we find that different *orderings* have different effects on the behaviour of the Wright-Fisher model for the population dynamics at two linked loci only, which is mainly caused by different effects of genetic recombination on natural selection for different *orderings*. More specifically, genetic recombination enhances the effect of natural selection when natural selection occurs before population regulation under the Wright-Fisher model. The effect of the *ordering* on the behaviour of the Wright-Fisher model gradually weakens as the population approaches linkage equilibrium and completely vanishes once linkage equilibrium has been reached. In addition, such effect becomes more significant along with the increasing selection coefficient, recombination rate or population size, and the duration of the effect of the *ordering* on the behaviour of the Wright-Fisher model decreases as the recombination rate or population size increases.

### 3.2. Robustness of Monte Carlo Simulations

In this section, we perform a robustness analysis of Monte Carlo simulations carried out in Section 3.1. The purpose of this robustness analysis is to demonstrate that the statistical noise due to a finite number of realisations becomes very small as the number of realisations becomes large, so that the discrepancy in Monte Carlo simulation results from different *orderings* are not due to statistical noise and are indeed caused by the effect of the *ordering*.

According to Van der Vaart (2000), the rate of convergence for the empirical transition probabilities obtained from Monte Carlo simulations with respect to the Hellinger distance is of order 1/*M*^1/4^, where *M* is the number of independent realisations for the Monte Carlo simulation study. Combining with Le Cam and Yang (2000), we find that the rate of convergence for the Hellinger distance approximated by Monte Carlo simulations is of order *N*^1/2^/*M*^1/4^, which implies that in theory, the statistical noise can be reduced with the increase of the number of independent realisations.

Figures 12 and 13 illustrate the dynamics of the Hellinger distance 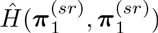 and the Hellinger distance 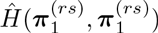, which are the Hellinger distance of the two empirical transition probabilities for the two runs of the same Wright-Fisher model, each simulating 10^5^ independent realisations, the *sr*-Wright-Fisher model in Figure 12 and the *rs*-Wright-Fisher model in Figure 13. In comparison with Figures 12 and 13, Figures 3 and 8 show that the statistical noise in the Monte Carlo approximation of the Hellinger distance is negligible, although there is some discrepancy for the extremely small selection coefficient in large populations. Obviously, this could be reduced further with more realisations. Hence, we conclude that the difference in the behaviour of the Wright-Fisher model between two different *orderings* indeed results from the effect of the *ordering*, rather than statistical noise.

**FIGURE 12.**
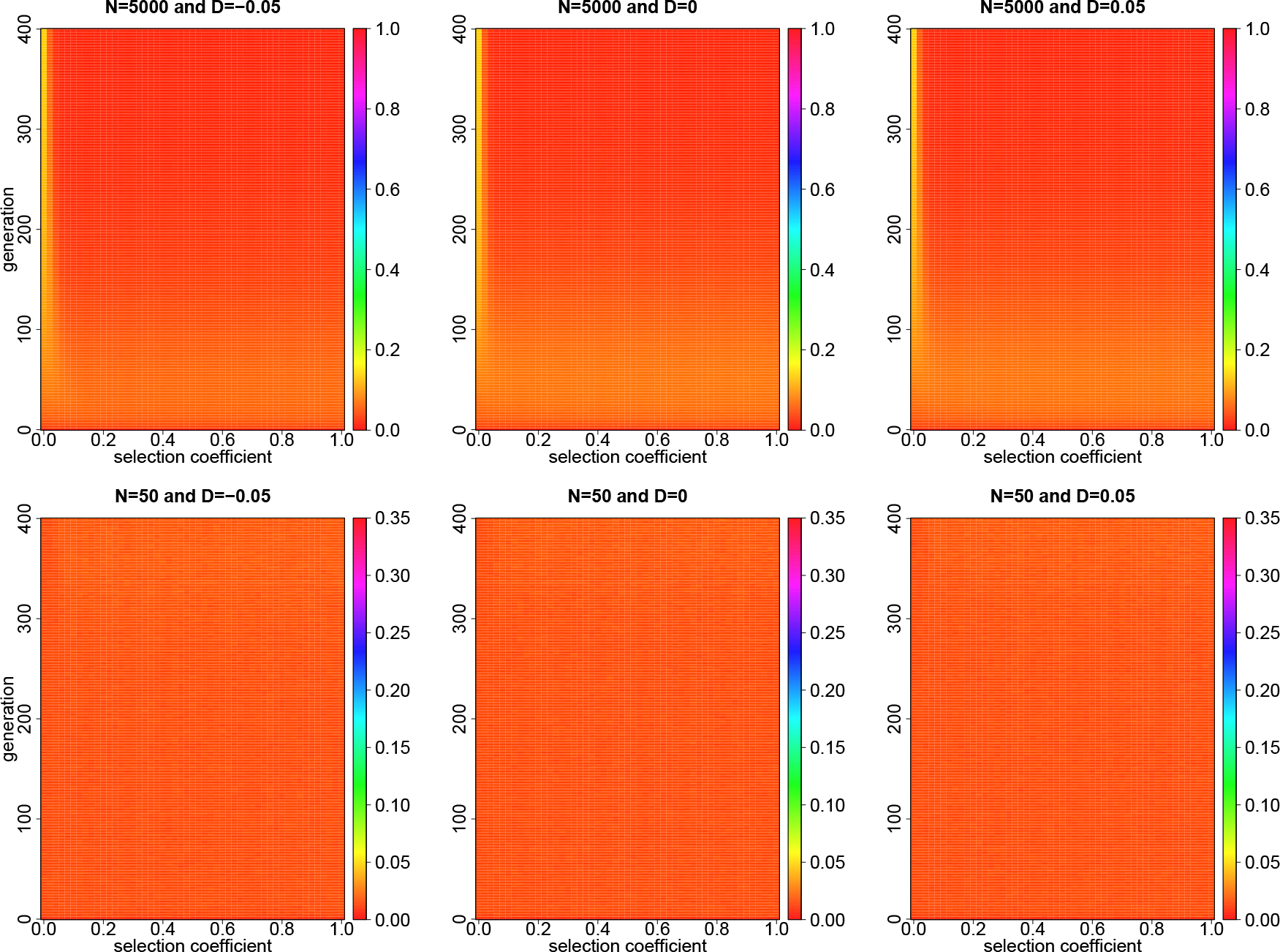
Dynamics of the Hellinger distance 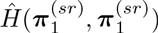 with varying selection coefficients 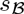 over 400 generations when the gene action at locus 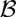 is completely additive. The Hellinger distance 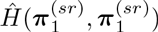 is evaluated from Monte Carlo simulations with 10^5^ independent realisations. The other parameters are 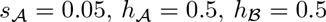 = 0.45.

**FIGURE 13.**
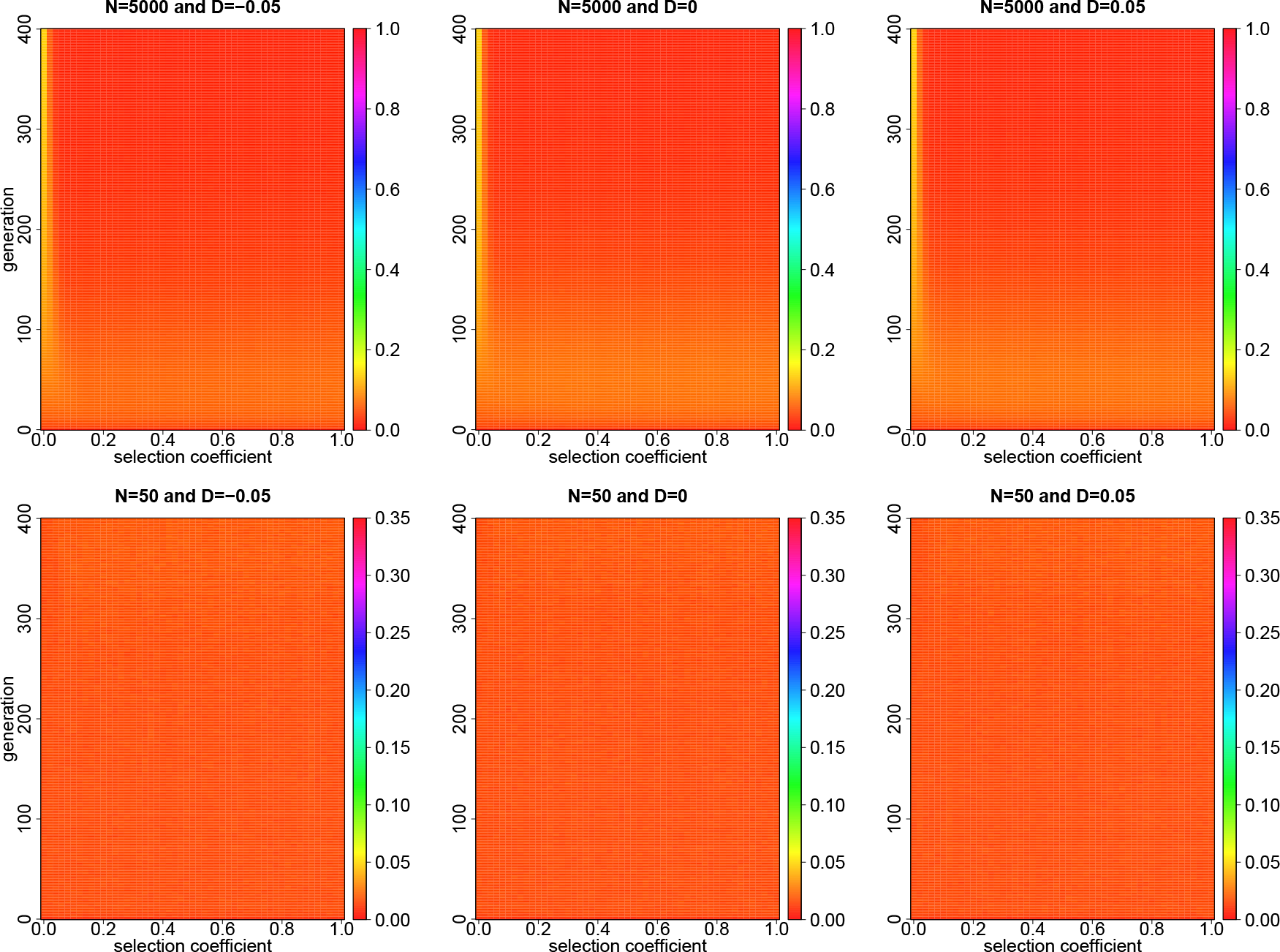
Dynamics of the Hellinger distance 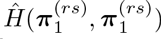 with varying selection coefficients 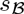 over 400 generations when the gene action at locus 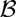 is completely additive. The Hellinger distance 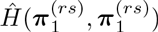 is evaluated from Monte Carlo simulations with 10^5^ independent realisations. The other parameters are 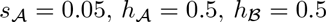 and *r* = 0.45.

## 4. Discussion and Conclusion

In this section, we investigate whether natural selection occurring before or after population regulation has different effects on the behaviour of the standard diffusion approximation of the Wright-Fisher model, which is commonly known as the Wright-Fisher diffusion. Moreover, we summarise our main results and end with some further perspectives.

### 4.1. Wright-Fisher Diffusion with Selection

The Wright-Fisher diffusion is one of the most important extensions of the Wright-Fisher model, which can be traced back to Kimura (1964). In recent years, the Wright-Fisher diffusion has already been successfully applied in the various population genetic analysis (*e.g*., Williamson et al., 2005; Bollback et al., 2008; Gutenkunst et al., 2009; Malaspinas et al., 2012; Steinrücken et al., 2014). More specifically, the Wright-Fisher diffusion is a limiting process of the Wright-Fisher model describing the changes in gamete frequencies over time in large populations under weak natural selection, which assumes that the selection coefficients (and recombination rates) are of order 1/(2*N*) and run time at rate 2*N*, *i.e*., *t* = *k*/(2*N*). Notice that we only present the formulation of the Wright-Fisher diffusion here and refer to Ethier and Nagylaki (1989) for a rigorous proof.

#### 4.1.1. *A Standard Diffusion Approximation.*

Let 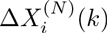 denote the change in the frequency of gamete *i* from generation *k* to the next, and then using Eq. (4), we have

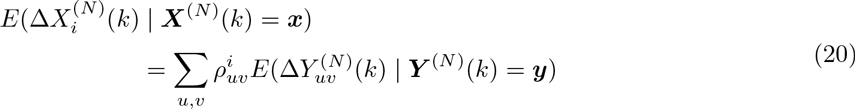

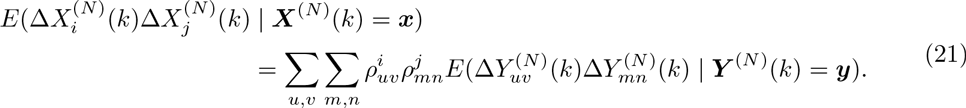

From Eq. (2), we have

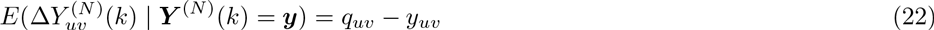

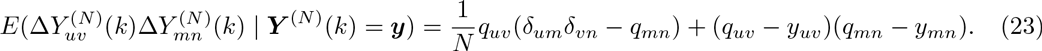

Substituting Eqs. (22) and (23) into Eqs. (20) and (21), respectively, we have

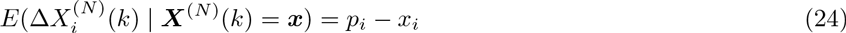

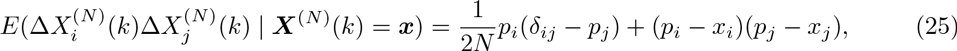

where

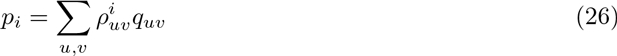

is the frequency of gamete *i* in an effectively infinite population after a single generation of the possible evolutionary forces (except genetic drift) at intermediate stages of the life cycle.

Considering the limits as the population size *N* goes to infinity, we can formulate the infinitesimal mean vector *μ*(*t, x*) as

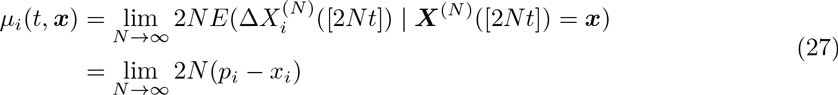

and the infinitesimal covariance matrix ***Σ***(*t*, *x*) as

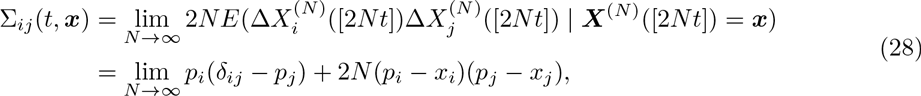

where [η] is used to denote the integer part of the value in the brackets, according to standard techniques of diffusion theory (see, for example, Durrett, 2008).

Therefore, the process of gamete frequencies ***X***^(*N*)^ converges to a diffusion process, denoted by ***X*** = {***X*** (*t*),*t* ≥ 0}, satisfying the following stochastic differential equation (SDE) of the form

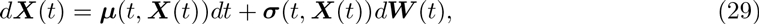

where the diffusion coefficient matrix *σ*(*x*) satisfies the relation that

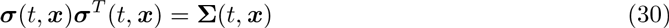

and ***W***(*t*) is a multi-dimensional standard Brownian motion. Here we refer to the process of gamete frequencies ***X*** = {***X***(*t*), *t* ≥ 0} as the Wright-Fisher diffusion with selection.

#### 4.1.2. *Effects of the* ordering *on the Wright-Fisher diffusion.*

Now we discuss whether different *orderings* have different effects on the behaviour of the Wright-Fisher diffusion, especially for the evolution of one- and two-locus systems under natural selection. As mentioned above, the Wright-Fisher diffusion is fully determined by the infinitesimal mean vector in Eq. (27) and the infinitesimal covariance matrix in Eq. (28). Our problem thereby reduces to checking whether the Wright-Fisher diffusion has the same SDE-representation for different *orderings*.

As demonstrated in Section 2.2, the one-locus system under natural selection has the same sampling probabilities ***q*** for the two *orderings*, *sr* and *rs*. Substituting Eq. (5) into (26), we obtain the same vector with frequencies of two possible alleles in an effectively infinite population after a single generation of natural selection *p* = (*p*_1_,*p*_2_) for the two *orderings, sr* and *rs*, and thereby the same SDE representation of the one-locus Wright-Fisher diffusion with selection. For the explicit expression of the one-locus Wright-Fisher diffusion driven SDE one is referred to Etheridge (2011). That is, the *ordering* does not affect the behaviour of the Wright-Fisher diffusion for the evolution of the one-locus system under natural selection.

However, as shown in Section 2.3, the two-locus system under natural selection has different sampling probabilities ***q*** for the two *orderings*, *sr* and *rs*, which implies that the vector with frequencies of four possible gametes in an effectively infinite population after a single generation of genetic recombination and natural selection *p* = (*p*_1_,*p*_2_,*p*_3_,*p*_4_) is different for the two *orderings*, *sr* and *rs.* Substituting Eqs. (12)-(13) and (14)-(15) into (26), respectively and using Taylor expansions with respect to the selection coefficients 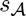 and 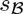, we have

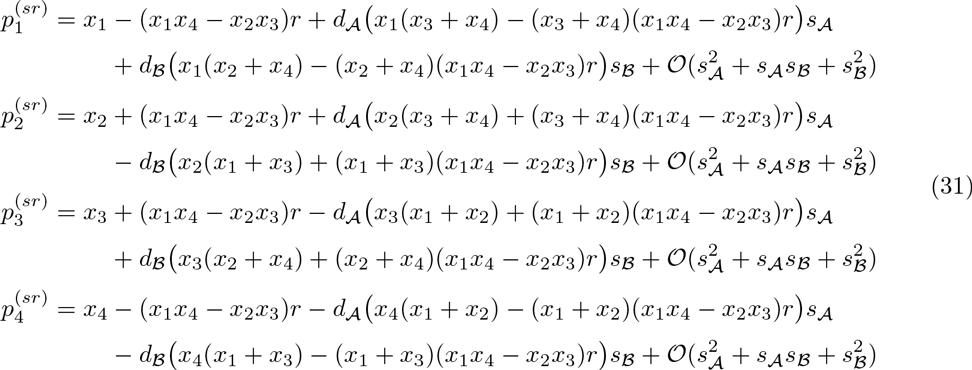

and

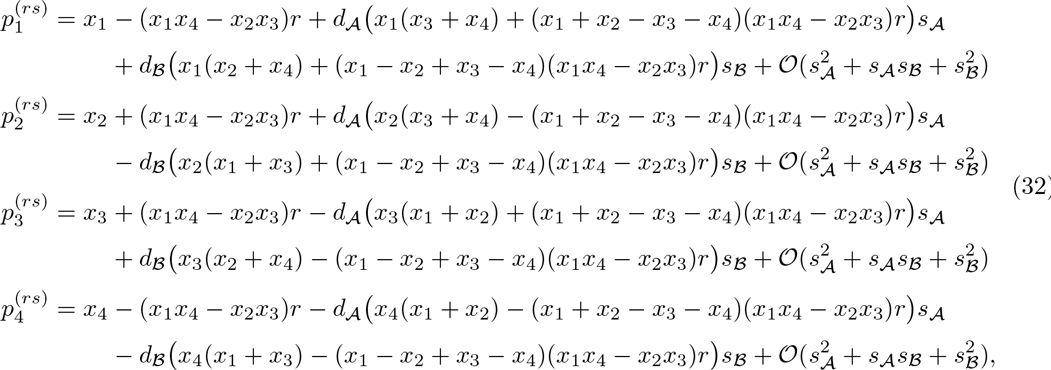

where

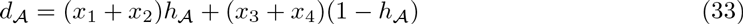

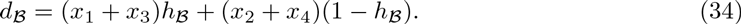

That is,we have

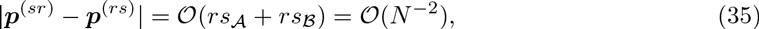

where | η | denotes the Euclidean vector norm. Substituting Eqs. (31) and (32) into (27) and (28), respectively, we have

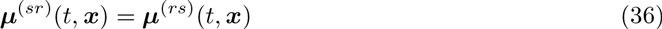

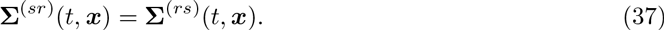

Eqs. (36) and (37) ensure the same SDE-representation of the two-locus Wright-Fisher diffusion with selection for the two *orderings*, *sr* and *rs.* That is, the *ordering* does not affect the behaviour of the Wright-Fisher diffusion for the evolution of the two-locus system under natural selection, which confirms that the difference caused by different *orderings* in the behaviour of the Wright-Fisher model is negligible for large populations evolving under weak natural selection at two tightly linked loci.

Therefore, we can conclude that the *ordering* does not affect the behaviour of the Wright-Fisher diffusion for the population dynamics at a single locus or two linked loci.

### 4.2. Summary and Further Perspectives

In this section, we summarise our main results obtained in the previous sections and discuss possible applications and extensions.

#### 4.2.1. *Summary.*

For diploid populations, natural selection occurring before or after population regulation only affects the behaviour of the Wright-Fisher model for the population dynamics at multiple linked loci, which is mainly caused by different effects of genetic recombination on natural selection for two different evolutionary sequences in the life cycle. More specifically, in the Wright-Fisher model, genetic recombination enhances the effect of natural selection when natural selection occurs before population regulation. The effect of natural selection occurring before or after population regulation on the behaviour of the Wright-Fisher model gradually weakens as the population approaches linkage equilibrium and completely vanishes once linkage equilibrium has been reached. Moreover, such effect becomes more significant as natural selection and genetic recombination become stronger or genetic drift becomes weaken. Strengthening the effect of genetic recombination or weakening the effect of genetic drift may reduce the duration of the effect of natural selection occurring before or after population regulation on the behaviour of the Wright-Fisher model. However, the effect of natural selection when natural selection occurs before population regulation completely vanishes in the Wright-Fisher diffusion for the population dynamics under natural selection.

For haploid populations, the study of how natural selection occurring before or after population regulation affects the behaviour of the Wright-Fisher model and its diffusion approximation follows in the similar manner to that employed here, and we can see that the results for haploid populations confirm that for diploid populations we have summarised above.

Given that most natural populations are probably near linkage equilibrium (Ridley, 2004), the different behaviour of the Wright-Fisher model caused by natural selection occurring before or after population regulation may not be significant. However, one evolutionary scenario in which it may have significant consequences is that of admixture between two populations. In this case, there is likely to be significant linkage disequilibrium due to allele frequency differences between populations (Pritchard and Rosenberg, 1999), and also, there is likely to be local selection in which alleles that are favoured in one population are selected against in the other (Charlesworth et al., 1997), and vice-versa. Therefore, the results presented here may lead to testable predictions about the role of life-history on the selection dynamics in natural populations. For example, we predict that recombination will have a more significant effect on the selection dynamics when natural selection occurs before population regulation.

#### 4.2.2. *Further Perspectives.*

In the present work, we only consider the evolutionary scenario that populations evolve under natural selection at fewer than two linked loci under the Wright-Fisher model, but the conditions required for this case are quite restrictive and probably not often met in natural populations. It would be interesting to take population structure, gene flow and mutation into account or consider the population dynamics at more than two loci, which provides opportunities to perform more reliable robustness analysis of the Wright-Fisher model with different possible sequences of the evolutionary processes in the life cycle. Furthermore, we investigate how natural selection occurring before or after population regulation affects the Wright-Fisher model through Monte Carlo simulations, which can only demonstrate the qualitative difference in the behaviour of the Wright-Fisher model. It would be far more challenging to carry out analysis of the effect of different sequences of the evolutionary processes in the life cycle on the Wright-Fisher model that can lead to more quantitative comparisons.

